# Programmed transport and release of nanoscale cargo by immune cells

**DOI:** 10.1101/846956

**Authors:** Daniel Meyer, Saba Telele, Anna Zelená, Elsa Neubert, Robert Nißler, Florian Mann, Luise Erpenbeck, Sarah Köster, Sebastian Kruss

**Affiliations:** Institute of Physical Chemistry, Göttingen University, 37077 Göttingen, Germany; Department of Dermatology, Venerology and Allergology, University Medical Center, Göttingen University, 37077 Göttingen, Germany; Institute of X-Ray Physics, Göttingen University, 37077 Göttingen, Germany

**Keywords:** carbon nanotubes, sensors/biosensors, nanobiotechnology, biomedical applications, photoluminescence

## Abstract

Transport and delivery of (nanoscale) materials are crucial for many applications in biomedicine. However, controlled uptake, transport and triggered release of such cargo remains challenging. In this study, we use human immune cells (neutrophilic granulocytes, neutrophils) and program them to perform these tasks in vitro. For this purpose, we let neutrophils phagocytose a nanoscale cargo. As an example, we used DNA-functionalized single-walled carbon nanotubes (SWCNT) that fluoresce in the near infrared (980 nm) and serve as sensors for small molecules. Cells still migrate, follow chemical gradients and respond to inflammatory signals after uptake of the cargo. To program release, we make use of neutrophil extracellular trap formation (NETosis), a novel cell death mechanism that leads to chromatin swelling and subsequent rupture of the cellular membrane and release of the cell’s whole content. By using the process of NETosis we can program the time point of cargo release *via* the initial concentration of stimuli such as phorbol 12-myristate-13-acetate (PMA) or lipopolysaccharide (LPS). At intermediate stimulation with LPS (100 μg/ml), cells continue to migrate, follow gradients and surface cues for around 30 minutes and up to several hundred micrometers until they stop and release their cargo. The transported and released SWCNT sensor cargo is still functional as shown by subsequent detection of the neurotransmitter dopamine and reactive oxygen species (H_2_O_2_). In summary, we hijack a biological process (NETosis) and demonstrate how neutrophils can be used for programmed transport and delivery of functional nanomaterials.

## 1. Introduction

Targeted delivery of (nano)materials and pharmaceuticals is one of the great challenges in biomedicine.^[1]^ Encapsulation of drugs in colloidal structures such as liposomes or polymeric micelles have been studied thoroughly over the years. These approaches demonstrated great success in delivery of active anti-cancer agents^[2–4]^, vitamins^[5]^, enzymes^[6,7]^ or antimicrobials.^[8]^ Nanomaterials such as nanoparticles^[9]^, carbon nanotubes^[10]^ or nanobots^[11]^ offer benefits due to their optoelectronic properties, tunable surface chemistry and ability to infiltrate cellular membranes.^[10]^ Additionally, they can be imparted with useful functions. For example, single-walled carbon nanotubes (SWCNTs) are known for their near infrared (nIR) fluorescence and serve as versatile building blocks for optical nanosensors.^[12–15]^ Their surface can be chemically tailored with DNA, peptides, lipids, nanobodies or viruses to sense biologically relevant signaling molecules with high spatiotemporal resolution.^[16–23]^ Thus, such nanomaterials are attractive candidates as cargo.

A large drawback of conventional drug delivery systems is their incapability to move autonomously. Most of the above mentioned approaches rely on an external flow (*e.g.* of the vascular system) to reach a target zone as they do not own a self-propelling mechanism. Therefore, crossing biological barriers and actively reaching a site of interest remains difficult.^[24,25]^

One way to overcome this issue is to equip the cargo transporter with additional capabilities. For example, magnetic nanoparticles have been manipulated through external magnetic fields.^[26]^ Another approach is to use and reprogram cells.^[27]^ Din *et al.* engineered bacteria for programmed lysis *in vivo* resulting in the delivery of cytotoxic agents and a potential way to tackle cancer.^[28]^ Another example is binding and transport of cargo molecules by surface-modified red blood cells, which form long-living, biocompatible hybrid carriers.^[29,30]^ Neutrophilic granulocytes (neutrophils) are the most abundant type of white blood cells. They are interesting candidates for cargo delivery because they are able to take up materials (phago-/endocytosis)^[31]^, sense and migrate along chemical gradients (chemotaxis) to inflammatory sites^[32,33]^ or cross dense borders, such as the blood-brain barrier.^[34,35]^ The abilities of neutrophils have been used to take up silica particles to follow *E.coli* gradients and paclitaxel containing liposomes for cancer treatment.^[34,36]^ However, so far it was not yet shown how to control cargo release. Another function of neutrophils is neutrophil extracellular trap (NET) formation (NETosis), a defense strategy and novel type of cell death.^[37,38]^ During NETosis, the cell’s chromatin is chemically modified, which leads to its expansion and ultimately the rupture of the cellular membrane and the release of their cytosolic content.^[39]^ Neubert *et al.* showed that this process consists of different phases, including a first active phase in which the cell remains fully functional.^[39]^ In summary, neutrophils possess several functions that are highly interesting for cargo delivery of nanoscale materials.

Here, we make use of these functions and demonstrate uptake, transport and programmed release of a nanoscale cargo. We show that neutrophils take up carbon-nanotube-based near infrared fluorescent sensors as cargo, transport them and release them again *via* NETosis. Importantly, we quantify time and length scales of this process and showcase that the cargo is fully functional after delivery by detecting small molecules with the transported nanosensors.

## 2. Results and Discussion

### 2.1 Uptake and release of carbon nanotubes by neutrophils

Neutrophils use phagocytosis to destroy foreign objects.^[31,40]^ Therefore, neutrophils should also be able to take up nanomaterials. In this study, we chose single-walled carbon nanotubes (SWCNT) as a model cargo to make use of their unique near infrared fluorescent sensing properties. Neutrophils readily took up DNA functionalized (GT)_15_-(6,5)-SWCNTs (**Fig.2a,b**). The nIR signal (red) of the uptaken SWCNTs is localized in a region of the neutrophil outside the nucleus (blue) and this compartment remained in the rear of the cells during migration (**Fig. 2a, Suppl. Movie 1**). It is most likely that this compartment is the phagosome, which is often found in the actomyosin-rich back of polarized neutrophils.^[41]^ Similarly, streptavidin-functionalized SWCNTs in HL60 cells, a model cell line for primary neutrophils were found in a similar location.^[42]^ The nuclei, on the other side, stayed rather at the middle/front of the cell during migration as previously described.^[43]^ Uptake of SWCNTs increased with concentration and incubation time as evidenced by the nIR fluorescent signal inside the cells (**Fig. 2b**). After around 15 min, uptake reached a plateau (**Fig. 2b**). For low SWCNT concentrations (0.1 nM) cells showed a normal behavior while at higher concentrations (> 1nM) we observed sometimes cell agglomerates (**Suppl. Fig. S1a**). For this reason, we used 0.1 nM SWCNT for uptake and all following experiments. Cells with a SWCNT cargo were still able to perform NETosis after stimulation with 100 nM PMA and demonstrated the well-documented time course of chromatin decondensation and subsequent cell rupture **(Fig. 2c)**. Interestingly, the distribution of the SWCNT cargo changed during NET-formation. The size of the intracellular SWCNT cargo did not change in early phases. In contrast, in later stages of NETosis the compartment with the SWCNTs were compressed, parallel to the expansion of the neutrophil’s chromatin (**Fig. 2c/d, Suppl. Fig. S2, Suppl. Movie 2**), which could be explained by an increasing intracellular pressure by chromatin swelling.^[39]^ This finding also explains why SWCNTs ended up in close proximity to the cell membrane.

Additionally, the SWCNT fluorescence intensity decreased during the time course of NETosis, which could be explained by changes in the pH or quenching because SWCNTs get closer to each other. Degradation of SWCNTs by myeloperoxidase (MPO) is well known but unlikely because it would occur on other time scales (hours to days).^[44,45]^

### 2.2 Functionality of cargo-loaded neutrophils

Uptake and release of the cargo is a necessary step for a delivery system. However, it remained elusive if DNA-SWCNT loaded cells activated to perform NETosis were still functional and able to migrate or for how long. Therefore, live-cell imaging of neutrophils exposed to different types and concentrations of NETosis inducing compounds (LPS and PMA) was performed. For this purpose, the random movement of the most motile cells (n = 30) for each condition and blood donor (N = 3 independent donors) were traced. Exemplary tracks are shown in **Fig. 3a** and trajectories for all conditions are shown in **Suppl. Fig.. S3**).

The time for the neutrophils to reach a stationary phase (stopping time) decreased with increasing concentration of the activator for both LPS and PMA as expected (**Fig. 3b**). The migration velocity stayed relatively constant for all LPS concentrations (**Fig. 3c**) but drastically decreased with increasing PMA concentration (≥10 nM). NETosis (indicated by chromatin decondensation) was assessed 160 min after activation and showed concentration dependent probabilities **(Fig. 3d**).

The results showed that low activator concentrations (0.1 – 10 μg/ml LPS and 0.1 - 1 nM PMA) did not trigger high NETosis rates but maintained the neutrophil’s motility. On the contrary, too high concentrations (10 –- 100 nM PMA) resulted in high decondensation rates but also inhibited cell migration completely. Only for medium concentrations (100 μg/ml LPS and 1 nM PMA), cells were still able to migrate and perform NETosis during the time course of the experiment (decondensation images are shown in **Suppl. Fig.. S4, example of cell migration behavior in Suppl. Movie 3**). In summary, we identified 100 μg/ml LPS as an optimal concentration to guarantee both migratory capabilities and efficient cargo release *via* NETosis.

### 2.3 Transport of nanoscale cargo via migration of neutrophils

To further investigate whether SWCNTs inside neutrophils affect their migration, a gradient migration assay (under-agarose migration) was performed with cargo loaded and unloaded cells.^[46]^ This assay mimics an *in vivo* scenario in which cargo loaded neutrophils are supposed to follow inflammatory cues and finally release their cargo at the inflammatory site. Both, experiments in commonly used fetal calve serum (FCS) environments (0.5%, **Fig. 4, Suppl. Fig.. S5**), as well as high concentrations (20%, **Suppl. Fig.. S6**), were performed and showed (**Suppl. Movie 4)** that cells react to chemokine gradients of interleukin-8 (IL8). In addition, cells were again exposed to different amounts of activators (0.1 – 10 nM PMA & 1 – 100 μg/ml LPS) and their migration distance was quantified after three hours of consecutive movement. In agreement with **Fig. 3**, increasing LPS concentrations decreased motility/covered distances. In contrast, addition of PMA showed either the same behavior as control samples (0.1 – 1 nM) or no migration at all (10 nM) implying an “all or nothing” behavior of PMA-induced NETosis pathways (**Fig. 4c**). This result is in agreement with a recent study that shows that PMA induced NETosis does not require adhesion or mechanical input at all.^[47]^ Interestingly, cells migrated over longer distances at higher serum concentration (20 % vs. 0.5 %) conditions, which highlights that this approach could also work *in vivo* (**Suppl. Fig. S5, S6** and **Suppl. Table T1/T2**). Cargo loaded and stimulated (LPS, 100 μg/ml) cells migrated between 145 ± 44 μm (0.5 % FCS) and 478 ± 162 μm (20 % FCS).

### 2.4 Programmed release of functional nanosensors

In the final step, we performed functionality tests of the SWCNT cargo in inactivated and ruptured cells to investigate whether the specific abilities of the internalized material remain intact throughout NETosis. SWCNTs are useful near infrared fluorescent building blocks of nanosensors for novel applications and their selectivity depends on the specific surface functionalization.^[13,48]^ We used (GT)_15_-(6,5) SWCNTs because they are able to report the presence of the neurotransmitter dopamine.^[16,49,50]^ Such sensors have been used to image dopamine release from cells with high spatiotemporal resolution.^[51]^ As a second cargo, hemin-aptamer functionalized SWCNTs that decrease their fluorescence in the presence of H_2_O_2_ were used.^[52,53]^ In both cases, SWCNTs were taken up by neutrophils and their responses were measured *via* consecutive nIR imaging either while they were carried by non-activated cells or after NETotic membrane rupture. Here, the addition of 100 nM dopamine led to an instantaneous increase of the sensors’ intensity in case of disrupted cells. In comparison, in non-damaged cells dopamine should not get into the cell and indeed such cells showed no sensor response to dopamine (**Fig. 5a, Suppl. Movie 5/6**). This result indicates a successful release and accessibility of the cargo after NETosis and full functionality of the dopamine nanosensors after release. In contrast, 100 μM H_2_O_2_ addition decreased the nIR signaling both intact and NETotic cells (**Fig. 5b**, **Suppl Movie 7/8**), which can be explained by the diffusion of H_2_O_2_ through the cellular membrane.^[54,55]^ Interestingly, we were also able to locate differences in the sensors’ uptake behavior depending on the associated surface functionalization. While (GT)_15_-(6,5) SWCNTs appeared most often in larger, intracellular structures, Aptamer/Hemin-SWCNTs were found closer to the cell membrane in smaller agglomerates (**Fig. 5a/b, Suppl. Fig. S7a**). Nevertheless, both sensor types were functional after cargo transport and rupture, as evidenced by the same fluorescence response performance prior to cellular uptake (**Suppl. Fig. S7b**) and control experiments (**Suppl. Fig. S7c/d**).

Finally, we also demonstrated the transport and release of the functionalized nanosensors to specific target sites. For this purpose, fibrinogen was patterned on glass surfaces and SWCNT-loaded, activated neutrophils were allowed to migrate over the coated area resulting in an alignment of the cells along the fibrinogen pattern (**Fig. 5c**). Again, the immobilized sensors were still functional after cell rupture on the pattern and showed an instant response to 100 nM dopamine (**Fig. 5d, Suppl. Movie 9**). Thus, neutrophils can be programmed to take up such a nanoscale cargo, transport it to specific sites and release it in a functional state.

Of course SWCNTs can be different in length, functionalization and chirality. It is known that those properties affect uptake and retention in cells^[56]^. Therefore, one could envision to tailor the nanoscale cargo for specific uptake/retention kinetics. In this study, functionalized SWCNTs have been used as sensors for two important signaling molecules (dopamine, H_2_O_2_). However, sensors for many other interesting molecules have been developed and could be transported by this approach^[16,17,20–23,57]^. For example, recently a SWCNT-based sensor for another important neurotransmitter (serotonin) has been introduced^[18]^. Another application is to use them as mechanical sensors. In this context, SWCNTs have been used to study movements in cells and the extracellular space in brain tissue^[58,59]^. Therefore, the cargo transport approach presented in this work could also be extended to bring SWCNT sensors into specific *in vivo* locations and explore the local mechanical properties.

## 3. Conclusion

Over the years much effort has been put forward into developing biocompatible transport and drug delivery systems. Here, we demonstrated a novel approach that makes use of particular and unique functions of neutrophils including phagocytosis, migration and NET formation. As we show, precise chemical activation of neutrophils determines how long the cells migrate and the time point of cargo release. In this process, the internalized nanoscale cargo remains functional at all times and is protected from most extracellular influences. This new type of transport-and-release mechanism might be of great benefit for various biomedical applications as it combines the biocompatibility and targeting capabilities of cells and a tool to program the time scale of release. For example, one could envision isolation of neutrophils from a patient, loading with functional nanoparticles and reinjection for programmed release. At the same time, this work also emphasizes the utility of SWCNTs as versatile chemical sensors that can be transported inside cells and are highly stable. In conclusion, we present a concept for nanoscale cargo transport and delivery by programming immune cells *via* NETosis.

## 4. Experimental Section

### Cell isolation of human neutrophils

Neutrophils were isolated from human venous blood of healthy donors. The study itself was approved by the ethics committee of the university medical center in Göttingen and all donors fully consent after being informed about possible consequences of the procedure. Isolation of human neutrophils was performed according to a standard protocol.^[60]^ In brief, fresh blood of healthy donors was collected with S-Monovettes KE 7.5 ml (Sarstedt) and layered gently on top of a Histopaque 1119 solution (ratio 1:1). After a first centrifugation step (1100 × g for 21 minutes), the transparent third, as well as the pink fourth layer, were collected and washed with HBSS (w/o Ca^2+^/Mg^2+^, Thermo Fisher Scientific). Cells were then again centrifuged for 10 minutes at 400 × g and the resulting pellet was resuspended in HBSS before layered on top of a phosphate buffered percoll (GE Healthcare) gradient with concentrations of 85, 80, 75, 70 and 65%. After a third centrifugation (1100 × g, 22 minutes), neutrophils were extracted by collecting half of the 70% and 80% layer as well as the entire 75% layer and washed once with HBSS. The remaining cell pellet was then resuspended in 1 ml HBSS, counted and lastly diluted to the needed concentration of the experiment. As a culture medium, RPMI 1640 (Lonza) with the addition of 10 mM HEPES and 0.5% fetal calf serum (FCS, Sigma-Aldrich) was used if not stated otherwise. Cellular identity was furthermore confirmed by a standard cytospin assay (Cytospin 2 Zentrifuge, Shannon) and a subsequent Diff-Quick staining (Medion Diagnostics). Cell purity had to exceed a 95% threshold to be used for any experiment.

### SWCNT modification with ssDNA

Surface modification of single-walled carbon nanotubes (SWCNTs) was performed as described previously.^[49,61]^ Briefly, 125 μl ssDNA solution (2 mg/ml stock in phosphate buffered saline (PBS))(Sigma Aldrich) and 125 μl (6,5) chirality enriched SWCNTs (Sigma Aldrich, Product No. 773735) (2 mg/ml stock in PBS) were placed for tip sonication (15 min / 30% Amplitude, Fisher Scientific TM Model 120 Sonic Dismembrator) in an ice bath. The obtained suspension was centrifuged 2 × 30 min / RT / 16.100 × g, while the Aptamer-SWCNT solution was furthermore excluded from the excess ssDNA using a Vivaspin 500 MWCO-filter (100.000 Da cut off). The sequence of the hemin-binding aptamer was (5‘-AGT GTG AAA TAT CTA AAC TAA ATG TGG AGG GTG GGA CGG GAA GAA GTT TAT TTT TCA CAC T-3’).^2^

### SWCNT uptake

To increase uptake rates, around 400000 cells were resuspended in 200 μl RPMI 1640 medium in a standard 1.5 ml Eppendorf tube and mixed 1:1 with an SWCNT solution (0.2 nM if not stated otherwise), diluted in RPMI 1640 medium. Cells were then incubated at 37 °C, 5% CO_2_ for 20 minutes, centrifuged once at 600 × g (5 minutes) and washed extensively with RPMI 1640 medium before seeding on the desired substrate.

The uptake analysis, shown in **Fig. 1b**, was performed likewise. Neutrophils were incubated, placed in a commonly used μ-slide 8 well chamber slide (ibidi, 75000 cells per well and 200 μl RPMI 1640 medium) and fixated using 4% paraformaldehyde (PFA) (Sigma Aldrich, CAS 30525-89-4) for 1 hour after letting the cells adhere for another 30 minutes at 37 °C, 5% CO_2_. Fixed samples were then placed in a custom build near-infrared microscope composed of a Olympus BX-51 housing (Olympus), regular fluorescence (X-Cite Series 120 Q lamp, EXFO) and white light illumination (TH4-200 lamp, Olympus), a 561 nm laser (Jive 500, Cobolt) and two cameras (Zyla 4.2 sCMOS, Andor and Xeva-1.7-320, Xenics) with a 900 nm long pass filter (FEL0900, Thorlabs) in front to visualize SWCNT excitation as well as phase contrast and fluorescence in parallel. For imaging, a 20x objective (MPLFLN20X, Olympus) was chosen. For each condition, four images at each side of a well were recorded in phase contrast and nIR-mode (100 mW laser power, 500 ms exposure time) using the Zyla camera and saved in separate 16-bit files for subsequent analysis.

**Figure 1.**
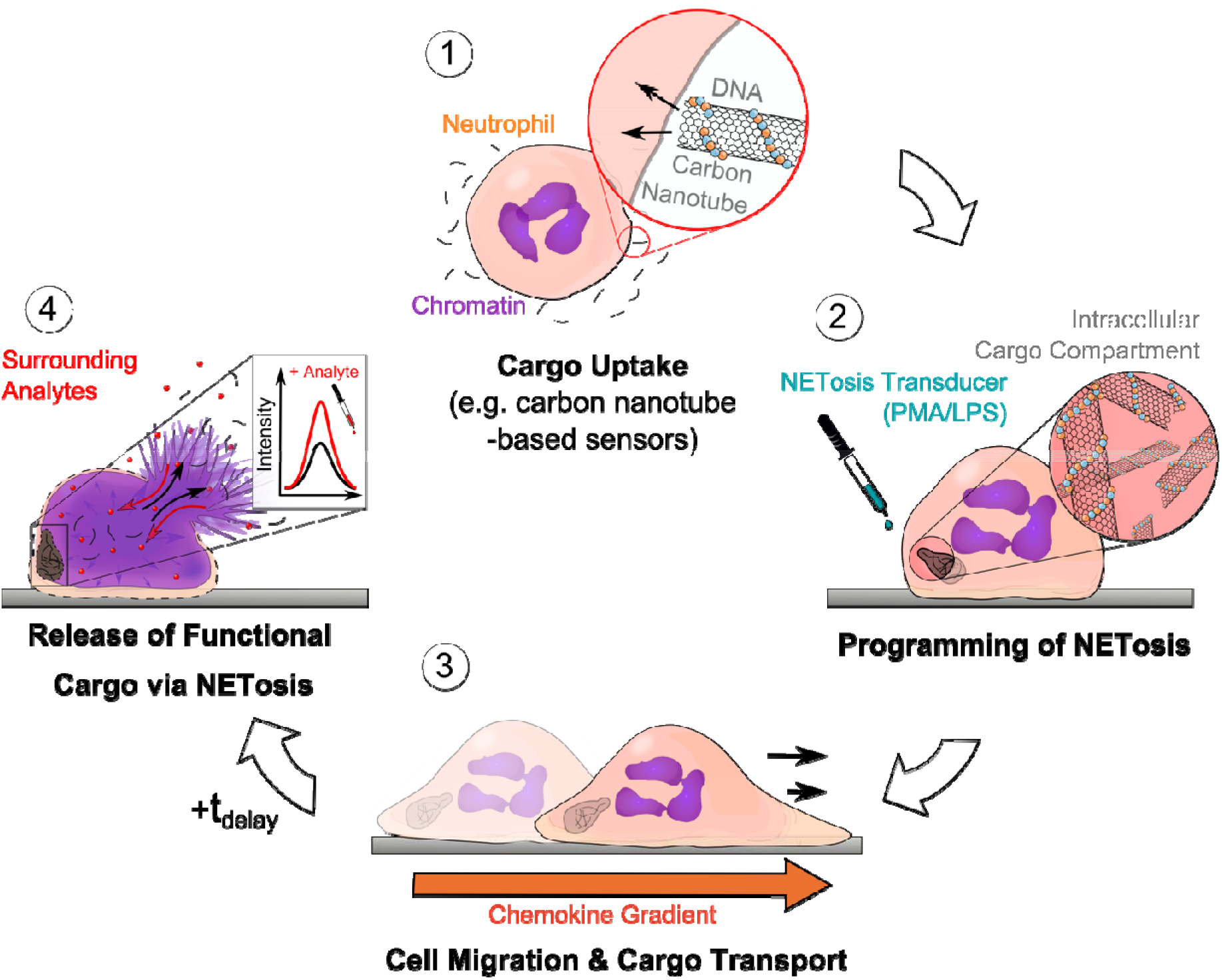
Schematic of uptake, transport and programmed release of cargo by neutrophils. (1) Neutrophils take up a nanomaterial such as DNA-coated single-walled carbon nanotube (SWCNT) sensors. (2) NETosis is chemically induced (with LPS or PMA), which determines the timepoint of NETosis and rupture. During the first phase after stimulation of NETosis cells are still functional and able to migrate and follow inflammatory signals (3). (4) Finally, at the end of the NETotic process, the cell membrane ruptures and releases the cargo into the extracellular space. In case of SWCNT sensors (as cargo) they sense analytes at the new location via their near infrared fluorescence signal.

**Figure 2.**
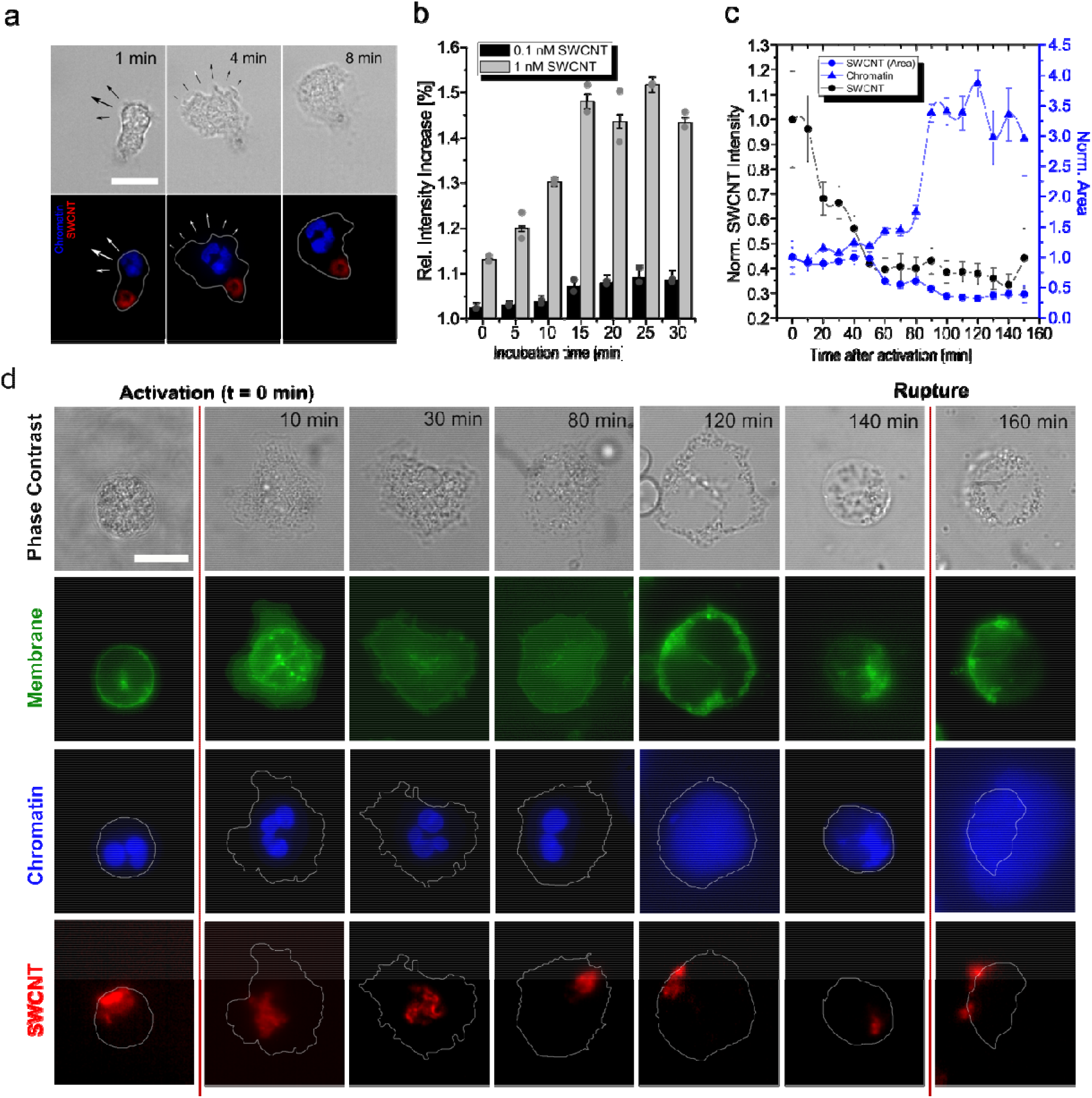
Uptake of nanomaterial cargo by neutrophils. **a** Neutrophils take up nanosensors (GT)_15_-SWCNT and are still fully functional and able to migrate. Phase contrast (top) and fluorescence images of chromatin (blue, Hoechst 33342 staining) and nIR fluorescent SWCNTs (red). SWCNTs appear to be located at the rear of the cell. Scale bar = 10 μm. **b** SWCNT uptake measured by nIR fluorescence intensity increase inside the cells. Uptake takes place within minutes and saturates after 15 minutes. Mean ± SEM, N = 2 donors, n > 60 cells for each time point. **c** The SWCNT fluorescence signal changes during NETosis. Both, SWCNT area (blue, circular data points) as well as intensity (black, circular data points) decrease during the process. Chromatin area (blue, triangular data points) increases as expected from NETosis. Mean ± SEM, N = 3 donors, n > 30 cells. **d** Time course of NETosis in different (GT)_15_-SWCNT loaded and activated neutrophils show cell rounding and chromatin decondensation. SWCNTs (> 900 nm, red), chromatin (blue), cell membrane (green), phase contrast (grey). SWCNTs appeared to be compressed/pushed by the expanding chromatin to the outer cell membrane in the final phase of NETosis. Scale bar = 10 μm.

**Figure 3.**
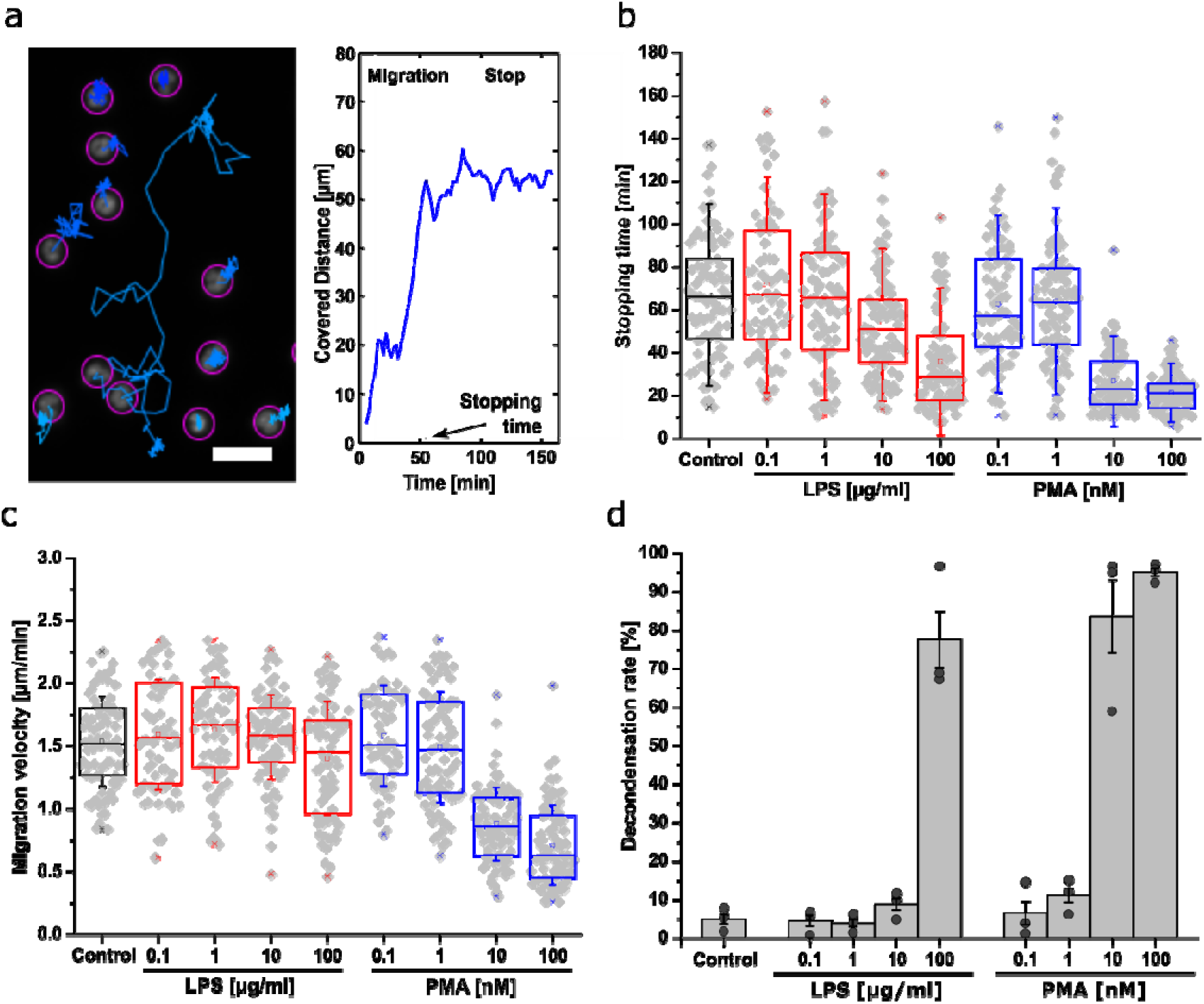
Migratory properties of cargo-loaded and NETosis programmed neutrophils. **<u>a</u>** <u>T</u>ypical trajectories (left) and the (absolute) covered distance of a neutrophil (right). Without inflammatory gradient cells randomly move in all directions until they stop due to the onset of the second phase of NETosis. Scale bar = 10 μm. **b-c** Time until the cells stop to move (stopping time) and migration velocity of activated neutrophils for different activation conditions. Increasing the concentration of LPS (red) decreases stopping time linearly. For PMA (blue) there was practically no movement above a certain concentration. Likewise, LPS does not influence the migration velocity whereas PMA slows them down at higher concentrations. N = 3 donors, n > 60 cells. Boxplot shows box line = 25-75% percentile, cross = mean, dot = min/max, error bars = SD. **d** Decondensation (NETosis) rates of neutrophils from **b**/**c** 160 minutes after activation. Very low amounts of LPS and PMA did not trigger NETosis. In contrast, higher values (100 μg/ml LPS, 10-100 nM PMA) lead to massive chromatin decondensation/NETosis. Data: mean ± SEM, N = 3, n > 60 cells. Nucleus stained with Hoechst 33342.

**Figure 4.**
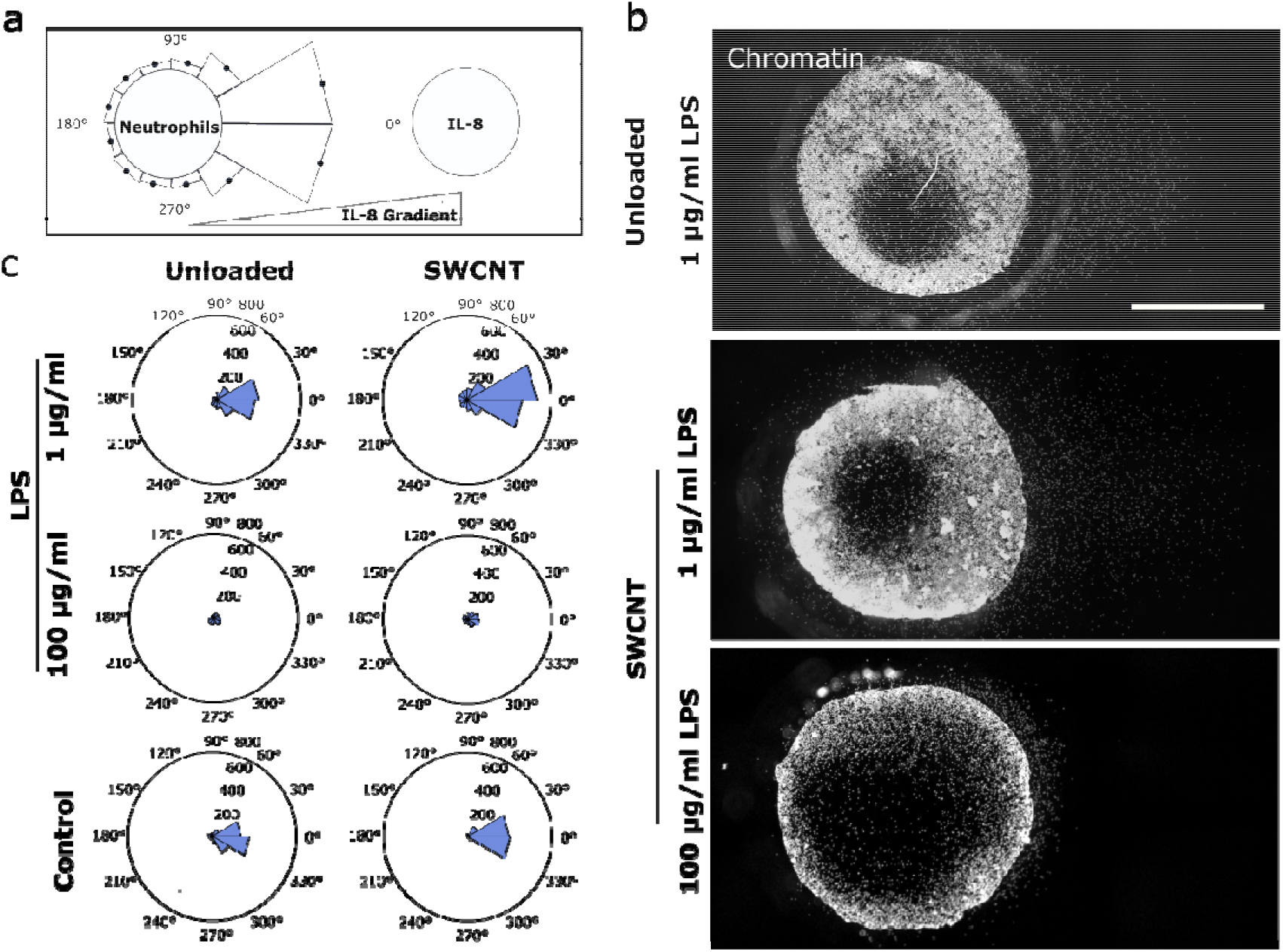
Collective nanoscale cargo transport by activated neutrophils. **a** Design of the migration experiment: Neutrophils programmed to perform NETosis migrate along an IL-8 gradient (under-agarose assay). Then the distance to the frontier of the leading cells is quantified. **b** The concentration of the NETosis activator affects the period during which the cells still migrate and the onset of NETosis. Images show nuclei of neutrophils (chromatin stained by Hoechst 33342) after 3h of migration. Scale bar = 500 μm. **c** Mean radial migration plots for different activators of NETosis and SWCNT uptake. Higher activator concentrations reduce the distance migrated by the cells. Interestingly, SWCNT-loaded cells travel around 30% further than the control cells after 3h.

**Figure 5.**
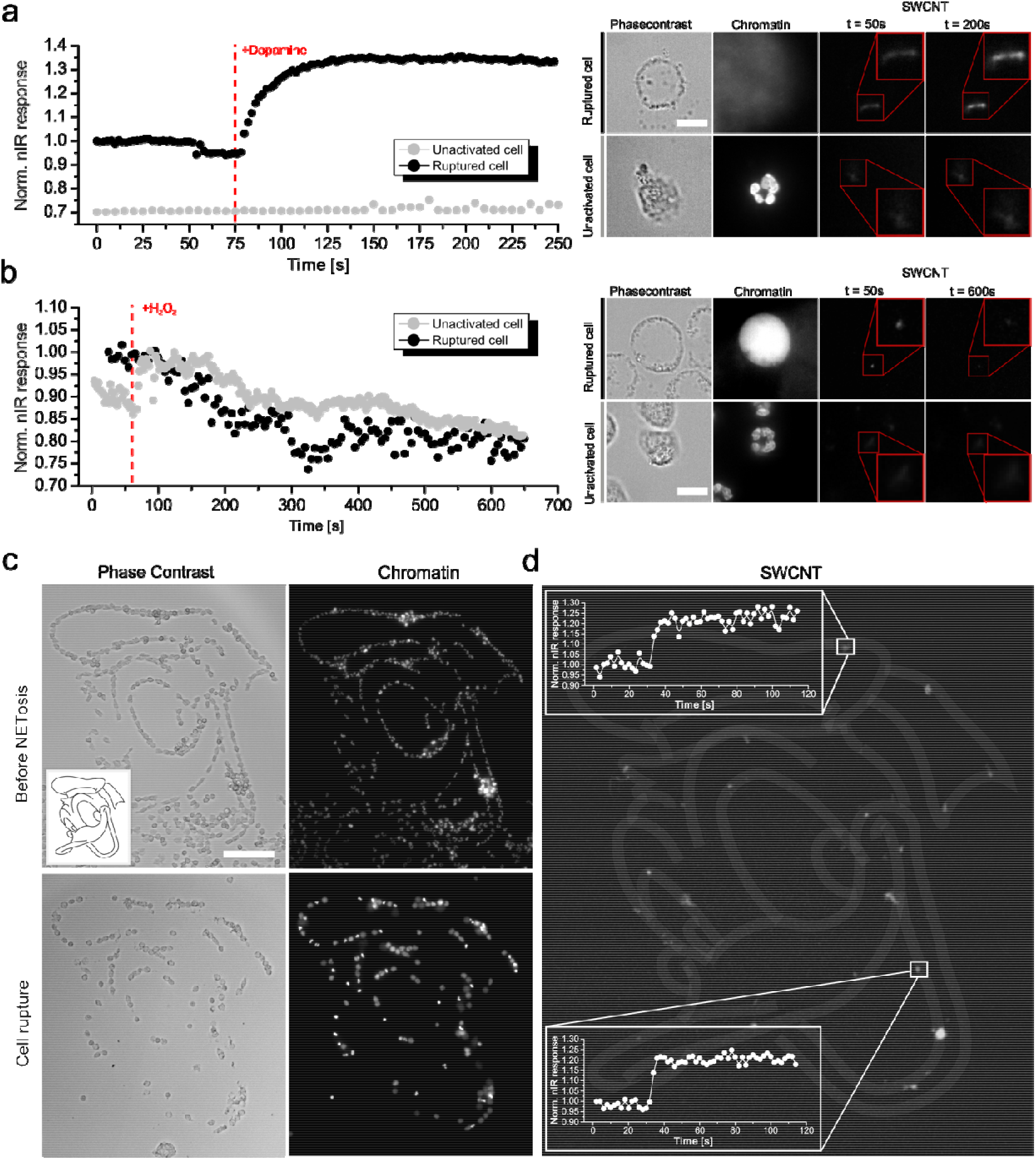
Release of functional cargo at target sites. **a** Cells programmed for NETosis release functional cargo such as (GT)_15_-(6,5) SWCNTs that serve as nIR sensors for the neurotransmitter dopamine (by intensity increase). SWCNTs inside non-activated cells (grey line) showed no fluorescence changes to 100 nM dopamine. In contrast, the area around cells that underwent NETosis showed a sensor response (black line). Scale bar = 5 μm, Chromatin stained with Hoechst 33342. **b** Other cargo such as Aptamer/Hemin-SWCNTs (H_2_0_2_ sensor) responded (intensity decrease) to 100 μM H_2_0_2_ both in inactivated (grey) and ruptured cells because H_2_0_2_ diffuses through cells membranes. **c** (GT)_15_-(6,5) SWCNT loaded neutrophils find adhesive cues and adhere to them. Here, fibrinogen was patterned on glass (top). After NETosis, chromatin (and cargo) is found in those locations (bottom). **d** nIR-colocalization of the (GT)_15_-(6,5) SWCNTs transported by neutrophils and the adhesive cues. Sensor positions correlated with the pattern (white lines) but the soft chromatin (including the sensors) quickly diffuses away and therefore the pattern does not remain stable. Sensors were still functional and respond to dopamine (100 nM). Scale bar = 100 μm. Chromatin stained with Hoechst 33342.

Measuring the intensity of SWCNT inside the cells was then performed using ImageJ’s thresholding system (v3.52i). nIR images were tuned to 8-bit depths, cropped into 20×20 μm areas containing only the respective SWCNTs and thresholded *via* common MinError thresholding algorithm to calculate a mean intensity. SWCNT spots that couldn’t be correlated with a cell area in the corresponding phase contrast image, as well as sensors that were found in cell agglomerations, were excluded from the analysis. The remaining data was averaged (weighted mean of all experiments) and normalized over the corresponding background value (I_br_ = 1) to calculate a normalized mean intensity for each condition.

### Live cell imaging of SWCNTs during NETosis

1 ml of RPMI medium containing 200000 (GT)_15_-(6,5) SWCNTs loaded cells and a 1 μg/ml concentration of Hoechst 33342 (Cat. H1399, Thermo Fisher Scientific) was poured on a glass bottom Petri dish (Cat. 150680, Thermo Fisher Scientific) and incubated to enable sufficient cell adhesion.

Subsequently, the sample was placed into a preheated incubation system (37 °C) (Cat. 11922, ibidi), on top of the custom build near-infrared microscope mentioned in the previous section. Using a 100x oil objective (UPLSAPO 100XO, Olympus) consecutively, phase contrast as well as chromatin (DAPI) and nIR-images were taken manually from the chosen sample position every ten minutes for 150 minutes after addition of 100nM phorbol myristate acetate (PMA) (Sigma-Aldrich) to the cell sample. SWCNTs were excited by a common fluorescence lamp in combination with a built-in 561nm filter cube (F48-553, AHF Analysentechnik) and excitation powers and exposure times were kept constant to ensure data comparability Analysis of chromatin and SWCNT area as well as SWCNT intensity was similar to the uptake study described before.

### Live cell imaging of activated neutrophil migration

To record the migration behavior of activated neutrophils, around 200 μl of RPMI 1640 medium containing 75000 untreated cells and 1μg/ml Hoechst 33342 stain were incubated in 8 Well μ-slides (ibidi) for 20 minutes Subsequently, the μ-slides were incorporated in pre-heated ibidi heating chambers (37 °C, 5% CO_2_, 90% humidity) on top of Olympus IX-81 microscopes. Using integrated XY-stages, 20x objectives (UCPLFLN 20x, Olympus) and corresponding phase contrast and DAPI channels (CBH white light lamp, U-HGLGPS fluorescence lamp, Olympus and 86-370-OLY DAPI filter-cube), a suitable position within each well was chosen and saved using the implemented steering software (Cell Sense, v. 2.1, Olympus). Subsequently, 200 μl of RPMI medium containing a distinct concentration of PMA (0.2, 2, 20, 200 nM) or lipopolysaccharide from *Pseudomonas Aeruginosa* (LPS) (0.2, 2, 20, 200 μg/ml) (Sigma-Aldrich) were added to a well in a blinded fashion and phase contrast as well as DAPI channels were recorded every two minutes for 160 minutes to image the cell movement.

### Migration analysis of preactivated neutrophils

Cell tracking was performed by ImageJ’s TrackMate plugin.^[62]^ Briefly, all chromatin images gained by the image process were combined to form a z-stack and implemented into the plugin first. Segmentation was then performed using the Laplacian of Gaussian (LoG) detector module with an estimated blob diameter of 20 pixels (6.5 μm) and a 5 pixel (1.6 μm) threshold. As a result, all cell trajectories within a stack could be traced back by using a subsequent simple LAP tracker model with a maximal linking and gap-closing distance of 50 pixels (16 μm) and a maximal frame gap of two frames. All trajectories were then further analyzed by a custom-build MATLAB code (v. Matlab 2014a) which was able to calculate the traveled distance *d* for every cell and frame *i* according to the formula

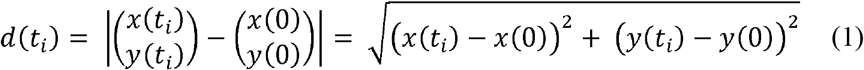

Subsequently, for each condition, 30 cells with the highest average distance values of the data set were further analyzed to detect the migration behavior of the most motile neutrophils within the environment. The stopping time of the migratory phase was calculated by plotting the distance-plots, likewise in a blinded fashion, together with the respective cell trajectory and manually searching for a time point of stagnation within the data set for each cell. Cells that did not show a clear change of moving patterns were excluded from the data set. The cell velocity of the migratory phase was then calculated by averaging the cell speed

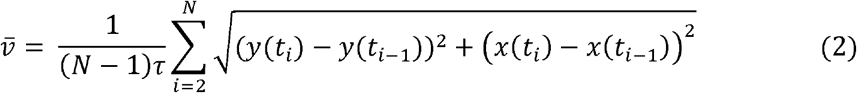

with *N* defining the number of frames until the stopping time and τ the frame time (2 minutes) between two images.

### Decondensation rate analysis

Counting decondensed and intact/lobular shaped nuclei was performed according to existing protocols.^[39,63]^ Briefly, chromatin images of the recorded positions were taken after 180 minutes and analyzed *via* ImageJ’s Cell Counter plugin. Nuclei that appeared in its known, compressed shape were counted and defined as intact/condensed whereas nuclei that showed increased, roundish chromatin distributions defined a basis for the decondensed/NETotic state. Cells were counted and the number of decondensed cells was divided by the total number of cells to generate a relative decondensation value.

### Migration in a gradient

Under-agarose assays were performed to measure cell migration of SWCNT-loaded neutrophils in a gradient. Gels were manufactured according to the protocol of B. Heit *et al.*.^[46]^ A HBSS/RPMI 1640 solution was prepared by mixing 5 ml HBSS (w/o Ca^2+^, Mg^2+^, Thermo Fisher Scientific) and 10 ml RPMI containing 0.75 % FCS (Merck) in a common 50-ml Eppendorf tube and heated up to 68°C using a common water bath. Meanwhile, 0.24 g ultra-pure agarose (Roth) was added to a vial containing 5 ml of milliQ and the solution was vortexed extensively in order to suspend the agarose homogeneously. The latter was subsequently heated up until boiling by the use of a common Bunsen burner and quickly vortexed for three times. The HBSS/RPMI solution was added to the agarose and 3 ml each of the mixture was evenly distributed on a plastic-bottom petri dish (Cat. 81156, ibidi). Agarose gels were then allowed to solidify at room temperature and samples were stored overnight at 4 °C with the dish lid covered in dust-free, milliQ saturated cloths to avoid gel draining. Shortly before the cell experiment, two wells with a diameter of 3 mm and a distance of 2.2 mm were punched in each gel using a dermal biopsy punch (Cat. KBP-48101, kai medical) and remaining agarose within each of the wells was extensively removed by vacuum aspiration. Lastly, gels were equilibrated with RPMI 1640 medium for one hour (37 °C, 5% CO_2_) and the supernatant medium was again removed by vacuum aspiration shortly before cell loading.

For the latter, around 100000 neutrophils ((GT)_15_-(6,5) SWCNTs loaded or without any pre-treatment) were poured in one of the prepared agarose wells using a 5 μl of medium plus 1.6 μM Hoechst 33342 solution and were allowed to equilibrate for 20 minutes at 37 °C, 5% CO_2_. Subsequently, 10 μl of a 0.1 μM IL-8 solution was poured into the second well while 5 μl of the respective NETosis activator concentration (LPS: 200, 20, 2 μg/ml, PMA: 20, 2, 0.2 nM or medium as control) was added to the cell well in the same time. All samples were stored for three hours in an incubator (37 °C, 5% CO_2_) and imaged afterward using the aforementioned IX-81 microscopic setup in combination with a 4x objective (UPLFLN 4X, Olympus). Afterward, cells were fixated by adding 2% PFA in PBS after carefully removing the gels with a scalpel and tweezers, washed twice with PBS, stored at 4 °C overnight and nIR-imaged by the aid of our aforementioned, customized nIR-setup the next day.

For image analysis, single images of each sample were stitched using ImageJ’s plugin MosaicJ. Fused images were background corrected and the existing cell distributions outside of the well were divided radially into 30-degree sections (see **Suppl. Fig. S5**) in which the distance of the leading cell edge was manually analyzed using ImageJ’s line tool. Resulting distances were averaged over all experiments and plotted as radial bar plots shown in **Fig. 4** or **Suppl. Fig. S4/5**or listed in **Suppl. Table T1/2**.

### nIR imaging of sensors

Cells for SWCNT functionality tests were prepared equally to those used for live cell imaging. Likewise, recordings of the sensor response in intact or ruptured neutrophils were performed at the customized build nIR-setup mentioned above in combination with an integrated ibidi heating chamber (37 °C) and the 100x oil objective. SWCNT of each condition were excited by a 561nm laser (200 mW) and recorded with a frame time of 2 or 5 seconds per image for four (dopamine addendum) or respectively 10 minutes (H_2_O_2_ addendum) in which, around the 1 minute mark, 500 μl of the particular analyte (300 nM dopamine, 300 μM H_2_O_2_, both in medium) was added. Subsequent exportation of image files, intensity analysis of single SWCNT areas or data handling was performed as described in the live cell imaging section.

### Cell patterning

Cell patterns on substrates shown in **Fig, 5c** were achieved by light-induced fibrinogen printing controlled by a PRIMO micropatterning machine (alvéole). Briefly, 18 × 18 mm glass coverslips (Fisher Scientific) were washed with 75% ethanol twice and plasma treated for five minutes to clean and improve the hydrophilicity of the sample. Subsequently, PDMS stencils with a circular well (*r* = 2 mm) in the center were pressed onto the glass and filled with 0.1 mg/ml PLL-g-PEG (Sigma) in PBS for one hour to guarantee a homogeneous passivation layer. Next, the so prepared sample was fixed on the respective PRIMO setup, calibrated according to the manufacturers’ instructions (microscope specs: Olympus IX83 with an UCPlanFL N 20× objective and an IX3-SSU stage) and filled with 5 μl of the photoactive reagent PLPP (1×, alvéole). Local radiation of the PLPP bound PEG-substrate using the patterning software LEONARDO (alvéole) then led to a protein degradation within the illuminated spots and enabled deposition of a second substrate protein. Sample were washed three times with PBS, coated again with Alexa-488 labeled fibrinogen solution (10 μl of a 50 μg/ml solution in PBS, Sigma) for twenty minutes and stored at 4 °C after another three washing steps. Patterns were normally used the next day, however, pattern degradation was not visible within one week after printing.

## Supporting Information

**Suppl. Movie 1** (AVI): SWCNT loaded neutrophils migrate in a 100 nM fMLP treated medium

**Suppl. Movie 2** (AVI): Multichannel live-cell images of PMA activated neutrophils (Phase Contrast, Chromatin, (GT)_15_-SWCNT)

**Suppl. Movie 3** (AVI): Migration and Decondensation behavior of differently activated neutrophils. Video shows Hoechst 33342 stained nuclei of unactivated cells (left), activated with 100 nM PMA (middle) or 100 μg/ml LPS (right).

**Suppl. Movie 4** (AVI): Neutrophils under agarose migration in an 1 μg/ml LPS medium

**Suppl. Movie 5** (AVI): nIR response of (GT)_15_-SWCNT loaded in a neutrophil during addition of 100 nM dopamine (no cell activation, cell & chromatin geometry shown in **Suppl. Fig. 7a**, grey dot marks timepoint of addition)

**Suppl. Movie 6** (AVI): nIR response of (GT)_15_-SWCNT loaded in a neutrophil during addition of 100 nM dopamine (after rupture, cell & chromatin geometry shown in **Suppl. Fig. 7a**, grey dot marks timepoint of addition)

**Suppl. Movie 7** (AVI): nIR response of aptamer coated Hemin-SWCNTs loaded in an unactivated, intact neutrophil during addition of 100 μM H_2_O_2_ (cell & chromatin geometry shown in **Suppl. Fig. 7a**, grey dot marks timepoint of addition)

**Suppl. Movie 8** (AVI): nIR response of Hemin coated-SWCNT loaded in a ruptured neutrophil during addition of 100 μM H_2_O_2_ (cell & chromatin geometry shown in **Suppl. Fig. 7a**, grey dot marks timepoint of addition)

**Suppl. Movie 9** (AVI): Activation of patterned SWCNTs with the aid of 100 nM dopamine addendum - grey dot marks timepoint of addition

## Acknowledgement

This project was supported by the state of Lower Saxony (life@nano) and the German Research Foundation (DFG grant KR 4242/4-1 and ER 723/2-1). We thank Andreas Janshoff and Claudia Steinem for fruitful discussions and support. We are grateful for fruitful discussions with members of the collaborative research center SFB 937 funded by the DFG. We thank the IMPRS PBCS for a PhD student fellowship (D.M.).

## References

[1] D. Peer, J. M. Karp, S. Hong, O. C. Farokhzad, R. Margalit, R. Langer, Nat. Nanotechnol. 2007, 2, 751.

[2] T. M. Allen, P. R. Cullis, Adv. Drug Deliv. Rev. 2013, 65, 36.

[3] A. H. Sarris, F. Hagemeister, J. Romaguera, M. A. Rodriguez, P. McLaughlin, A. M. Tsimberidou, L. J. Medeiros, B. Samuels, O. Pate, M. Oholendt, H. Kantarjian, C. Burge, F. Cabanillas, Ann. Oncol. Off. J. Eur. Soc. Med. Oncol. 2000, 11, 69.

[4] T. Nakanishi, S. Fukushima, K. Okamoto, M. Suzuki, Y. Matsumura, M. Yokoyama, T. Okano, Y. Sakurai, K. Kataoka, J. Control. Release 2001, 74, 295.

[5] M. Mohammadi, B. Ghanbarzadeh, H. Hamishehkar, Adv. Pharm. Bull. 2014, 4, 569.

[6] R. Ulbrich, R. Golbik, A. Schellenberger, Biotechnol. Bioeng. 1991, 37, 280.

[7] V. P. Torchilin, E. G. Tischenko, V. N. Smirnov, E. I. Chazov, J. Biomed. Mater. Res. 1977, 11, 223.

[8] S. Emami, S. Azadmard-Damirchi, S. H. Peighambardoust, H. Valizadeh, J. Hesari, J. Exp. Nanosci. 2016, 11, 737.

[9] L. R. Hirsch, R. J. Stafford, J. A. Bankson, S. R. Sershen, B. Rivera, R. E. Price, J. D. Hazle, N. J. Halas, J. L. West, Proc. Natl. Acad. Sci. U. S. A. 2003, 100, 13549.

[10] A. Bianco, K. Kostarelos, M. Prato, Curr. Opin. Chem. Biol. 2005, 9, 674.

[11] Y. Saadeh, D. Vyas, Am. J. Robot. Surg. 2014, 1, 4.

[12] M. J. Connell, S. M. Bachilo, C. B. Huffman, V. C. Moore, M. S. Strano, E. H. Haroz, K. L. Rialon, P. J. Boul, W. H. Noon, C. Kittrell, J. Ma, R. H. Hauge, R. B. Weisman, R. E. Smalley, Science. 2002, 297, 593 LP.

[13] S. Kruss, A. J. A. J. Hilmer, J. Zhang, N. F. N. F. Reuel, B. Mu, M. S. M. S. Strano, Adv. Drug Deliv. Rev. 2013, 65, 1933.

[14] E. Polo, S. Kruss, Anal. Bioanal. Chem. 2016, 408, 2727.

[15] G. Hong, S. Diao, A. L. Antaris, H. Dai, Chem. Rev. 2015, 115, 10816.

[16] F. A. F. A. Mann, N. Herrmann, D. Meyer, S. Kruss, Sensors (Switzerland) 2017, 17.

[17] J. D. Harvey, P. V. Jena, H. A. Baker, G. H. Zerze, R. M. Williams, T. V. Galassi, D. Roxbury, J. Mittal, D. A. Heller, Nat. Biomed. Eng. 2017, 1, 1.

[18] M. Dinarvand, E. Neubert, D. Meyer, G. Selvaggio, F. A. Mann, L. Erpenbeck, S. Kruss, Nano Lett. 2019.

[19] F. A. Mann, J. Horlebein, N. F. Meyer, D. Meyer, F. Thomas, S. Kruss, Chem. - A Eur. J. 2018, 24, 12241.

[20] G. Bisker, J. Dong, H. D. H. D. Park, N. M. N. M. Iverson, J. Ahn, J. T. J. T. Nelson, M. P. M. P. Landry, S. Kruss, M. S. M. S. Strano, Nat. Commun. 2016, 7, 1.

[21] A. A. Boghossian, B. Lambert, E. Ahunbay, V. Zubkovs, N. Schuergers, Small 2017, 13, 1701654.

[22] F. A. Mann, Z. Lv, J. Grosshans, F. Opazo, S. Kruss, Angew. Chemie Int. Ed. 2019.

[23] N. M. Bardhan, D. Ghosh, A. M. Belcher, Nat. Commun. 2014, 5, 1.

[24] A. C. Anselmo, J. Control. Release 2014, 190, 531.

[25] W. H. De Jong, P. J. A. Borm, Int. J. Nanomedicine 2008, 3, 133.

[26] C. C. Berry, S. Wells, S. Charles, A. S. G. Curtis, Biomaterials 2003, 24, 4551.

[27] V. Agrahari, V. Agrahari, A. K. Mitra, Expert Opin. Drug Deliv. 2017, 14, 285.

[28] M. O. Din, T. Danino, A. Prindle, M. Skalak, J. Selimkhanov, K. Allen, E. Julio, E. Atolia, L. S. Tsimring, S. N. Bhatia, J. Hasty, Nature 2016, 536, 81.

[29] C. H. Villa, A. C. Anselmo, S. Mitragotri, V. Muzykantov, Adv. Drug Deliv. Rev. 2016, 106, 88.

[30] J. Shi, L. Kundrat, N. Pishesha, A. Bilate, C. Theile, T. Maruyama, S. K. Dougan, H. L. Ploegh, H. F. Lodish, Proc. Natl. Acad. Sci. U. S. A. 2014, 111, 10131.

[31] H. H. Gustafson, D. Holt-Casper, D. W. Grainger, H. Ghandehari, Nano Today 2015, 10, 487.

[32] R. David, Nat. Rev. Mol. Cell Biol. 2013, 14, 547.

[33] F. Wang, Cold Spring Harb. Perspect. Biol. 2009, 1, a002980.

[34] C. Zhang, Z. Zhao, J. Xue, L. Wang, R. Mo, S. Shen, L. Xue, Z. Wei, L. Zhang, Y. Wen, H. Sun, Q. Ping, L. Kong, Nat. Nanotechnol. 2017, 12, 692.

[35] D. Chu, X. Dong, X. Shi, C. Zhang, Z. Wang, Adv. Mater. 2018, 30, 1706245.

[36] J. Shao, M. Xuan, H. Zhang, X. Lin, Z. Wu, Q. He, Angew. Chemie - Int. Ed. 2017, 56, 12935.

[37] B. V., R. U., G. C., F. B., U. Y., W. D.S., W. Y., Z. A., Science (80−.). 2004, 303, 1532.

[38] T. A. Fuchs, U. Abed, C. Goosmann, R. Hurwitz, I. Schulze, V. Wahn, Y. Weinrauch, V. Brinkmann, A. Zychlinsky, J. Cell Biol. 2007, 176, 231.

[39] E. Neubert, D. Meyer, F. Rocca, G. Günay, A. Kwaczala-Tessmann, J. Grandke, S. Senger-Sander, C. Geisler, A. Egner, M. P. Schön, L. Erpenbeck, S. Kruss, Nat. Commun. 2018, 9, 3767.

[40] C. Rosales, Front. Physiol. 2018, 9, 1.

[41] T. Mitchell, A. Lo, M. R. Logan, P. Lacy, G. Eitzen, Am. J. Physiol. Cell Physiol. 2008, 295, C1354.

[42] N. W. S. Kam, T. C. Jessop, P. A. Wender, H. Dai, J. Am. Chem. Soc. 2004.

[43] H. R. Manley, M. C. Keightley, G. J. Lieschke, Front. Immunol. 2018, 9, 2867.

[44] I. I. Vlasova, A. V. Sokolov, A. V. Chekanov, V. A. Kostevich, V. B. Vasilyev, Russ. J. Bioorganic Chem. 2011, 37, 453.

[45] K. A. Gogick, S. Petoud, W. A. Saidi, B. A. Barth, Y. Zhao, G. P. Kotchey, C. F. Chiu, A. Star, J. Am. Chem. Soc. 2013, 135, 13356.

[46] B. Heit, P. Kubes, Sci. Signal. 2003.

[47] L. Erpenbeck, A. L. Gruhn, G. Kudryasheva, G. Guenay, D. Meyer, E. Neubert, J. Grandke, M. P. Schoen, F. Rehfeldt, S. Kruss, bioRxiv 2019, 508366.

[48] J. P. Giraldo, H. Wu, G. M. Newkirk, S. Kruss, Nat. Nanotechnol. 2019, 14, 541.

[49] S. Kruss, M. P. Landry, E. Vander Ende, B. M. A. Lima, N. F. Reuel, J. Zhang, J. Nelson, B. Mu, A. Hilmer, M. Strano, J. Am. Chem. Soc. 2014, 136, 713.

[50] E. Polo, S. Kruss, J. Phys. Chem. C 2016, 120.

[51] S. Kruss, D. P. Salem, L. Vuković, B. Lima, E. Vander Ende, E. S. Boyden, M. S. Strano, Proc. Natl. Acad. Sci. 2017, 114, 1789.

[52] J. Pan, H. Zhang, T.-G. G. Cha, H. Chen, J. H. Choi, T.-G. G. Cha, H. Zhang, J. Pan, H. Chen, Anal. Chem. 2013, 85, 8391.

[53] H. Atsumi, A. M. Belcher, ACS Nano 2018, 12, 7986.

[54] J.-H. Kim, C. R. Patra, J. R. Arkalgud, A. A. Boghossian, J. Zhang, J.-H. Han, N. F. Reuel, J.-H. Ahn, D. Mukhopadhyay, M. S. Strano, ACS Nano 2011, 5, 7848.

[55] Y. Ohno, J. I. Gallin, J. Biol. Chem. 1985.

[56] M. Gravely, M. M. Safaee, D. Roxbury, Nano Lett. 2019, 19, 6203.

[57] N. F. Reuel, B. Grassbaugh, S. Kruss, J. Z. Mundy, C. Opel, A. O. Ogunniyi, K. Egodage, R. Wahl, B. Helk, J. Zhang, Z. I. Kalcioglu, K. Tvrdy, D. O. Bellisario, B. Mu, S. S. Blake, K. J. Van Vliet, J. C. Love, K. D. Wittrup, M. S. Strano, ACS Nano 2013, 7, 7472.

[58] N. Fakhri, A. D. Wessel, C. Willms, M. Pasquali, D. R. Klopfenstein, F. C. MacKintosh, C. F. Schmidt, Science (80-.). 2014, 344, 1031.

[59] A. G. Godin, J. A. Varela, Z. Gao, N. Danné, J. P. Dupuis, B. Lounis, L. Groc, L. Cognet, Nat. Nanotechnol. 2017, 12, 238.

[60] A. Mantovani, M. A. Cassatella, C. Costantini, S. Jaillon, Nat. Rev. Immunol. 2011, 11, 519.

[61] R. Nißler, F. A. Mann, P. Chaturvedi, J. Horlebein, D. Meyer, L. Vuković, S. Kruss, J. Phys. Chem. C 2019, 123, 4837.

[62] J. Y. Tinevez, N. Perry, J. Schindelin, G. M. Hoopes, G. D. Reynolds, E. Laplantine, S. Y. Bednarek, S. L. Shorte, K. W. Eliceiri, Methods 2017.

[63] S. L. Wong, M. Demers, K. Martinod, M. Gallant, Y. Wang, A. B. Goldfine, C. R. Kahn, D. D. Wagner, Nat. Med. 2015.

