## Supplementary_information for "Programmed transport and release of nanoscale cargo by immune cells"

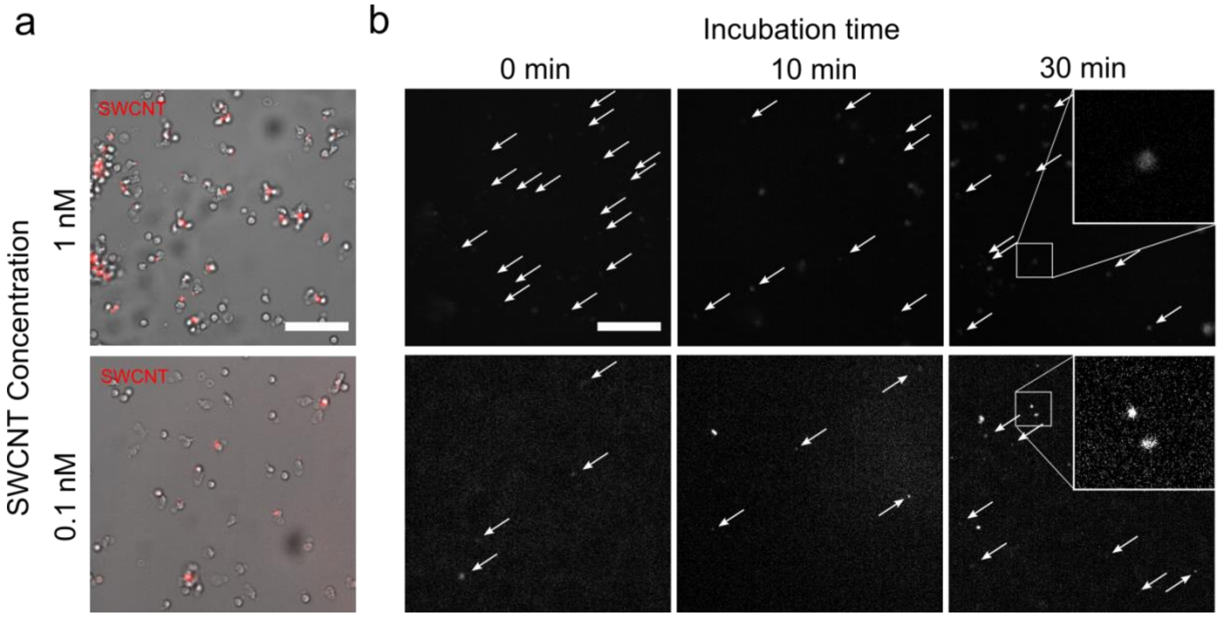

**Suppl. Fig. S1 SWCNT uptake by neutrophils. a** Phase contrast (grey) images of neutrophilic granulocytes after $(GT)_{15}$-SWCNT uptake. nIR SWCNT fluorescence (red) colocalized for higher concentrations(1 nM for 30 minutes, top) but there was also a tendency to cell aggregation. For lower SWCNT concentrations (0.1 nM, bottom) there was no cell aggregation but also less SWCNT signal. Normally, sensors could be found at the cell rear during migration, whereas in cell agglomerates SWCNT signals were located in the center of the cell bulk. Scale bar = 100 µm. **b** nIR images of uptaken $(GT)_{15}$-SWCNTs for different amounts of incubation time. SWCNT fluorescence signals increased with incubation time and starting concentration. Scale bar = 100 µm. The contrast of images at the bottom was increased to show SWCNT locations.

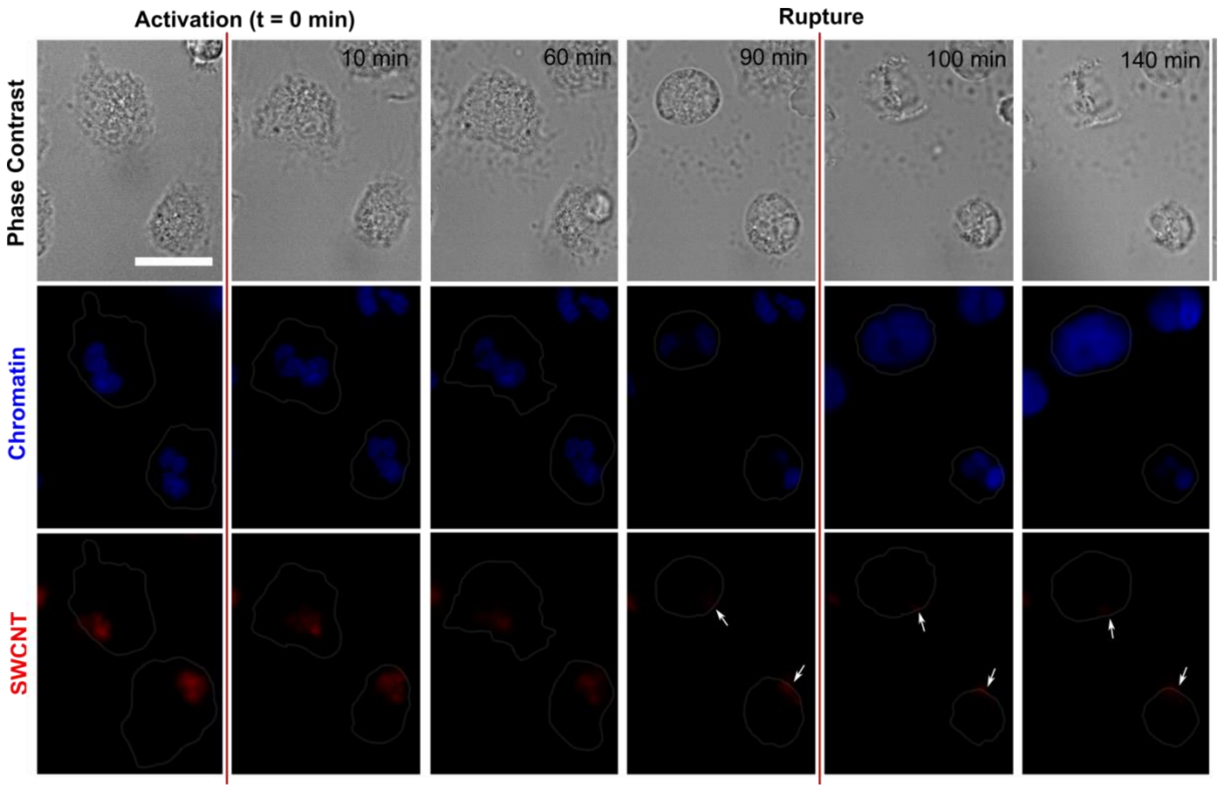

**Suppl. Fig. S2 SWCNT and chromatin geometry during NETosis.** Activated & SWCNT loaded cells show tight adhesion in the early phase of NETosis (phase contrast, top) while the uptaken SWCNTs (bot) and nuclei (mid) remain in their condensed compartments. In later stages, however, SWCNTs start to relocate to the cellular membrane resulting in a reduction of the sensor area and their overall fluorescence intensity. In the same time, the cells begin to deform while innercellular chromatin decondenses and mixes with the cytosolic content. Finally, in the last phase of NETosis, the cellular membrane ruptures and releases the decondensed chromatin as well as parts of the incorporated sensors. Scale bar = 10 µm, chromatin stained with Hoechst 33342.

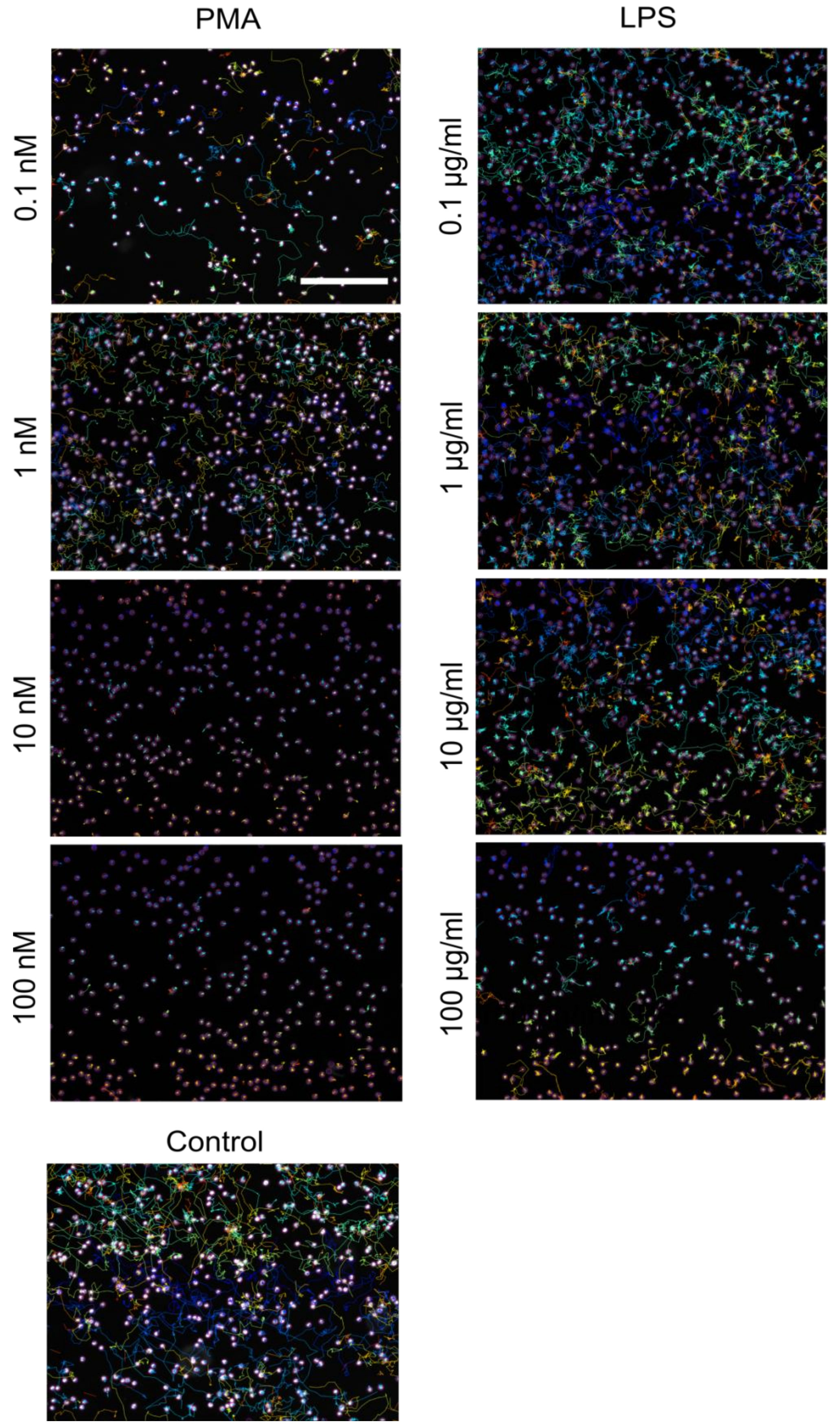

**Suppl. Fig. S3 Trajectories of neutrophils activated with different concentrations of NET-formation inducers.** PMA (0.1 – 100 nM) and LPS (0.1 – 100 µg/ml) stimulate different NET-formation pathways. Images show chromatin stained neutrophils and their trajectories. Low amounts of PMA led to similar speed and stopping time of migrating cells compared to control samples. On the contrary, 10 – 100 nM PMA forced nearly all cells to instantly stop. Lower amounts of LPS, did not change the movement pattern of the cells, however increasing concentrations lowered the cells speed and locomotion duration. Hoechst stain plus migration traces generated by TrackMate ImageJ plugin. Scale bar = 100 µm. Traces show migration patterns after/during 160 minutes. The track's color indicates only the cell index in the image.

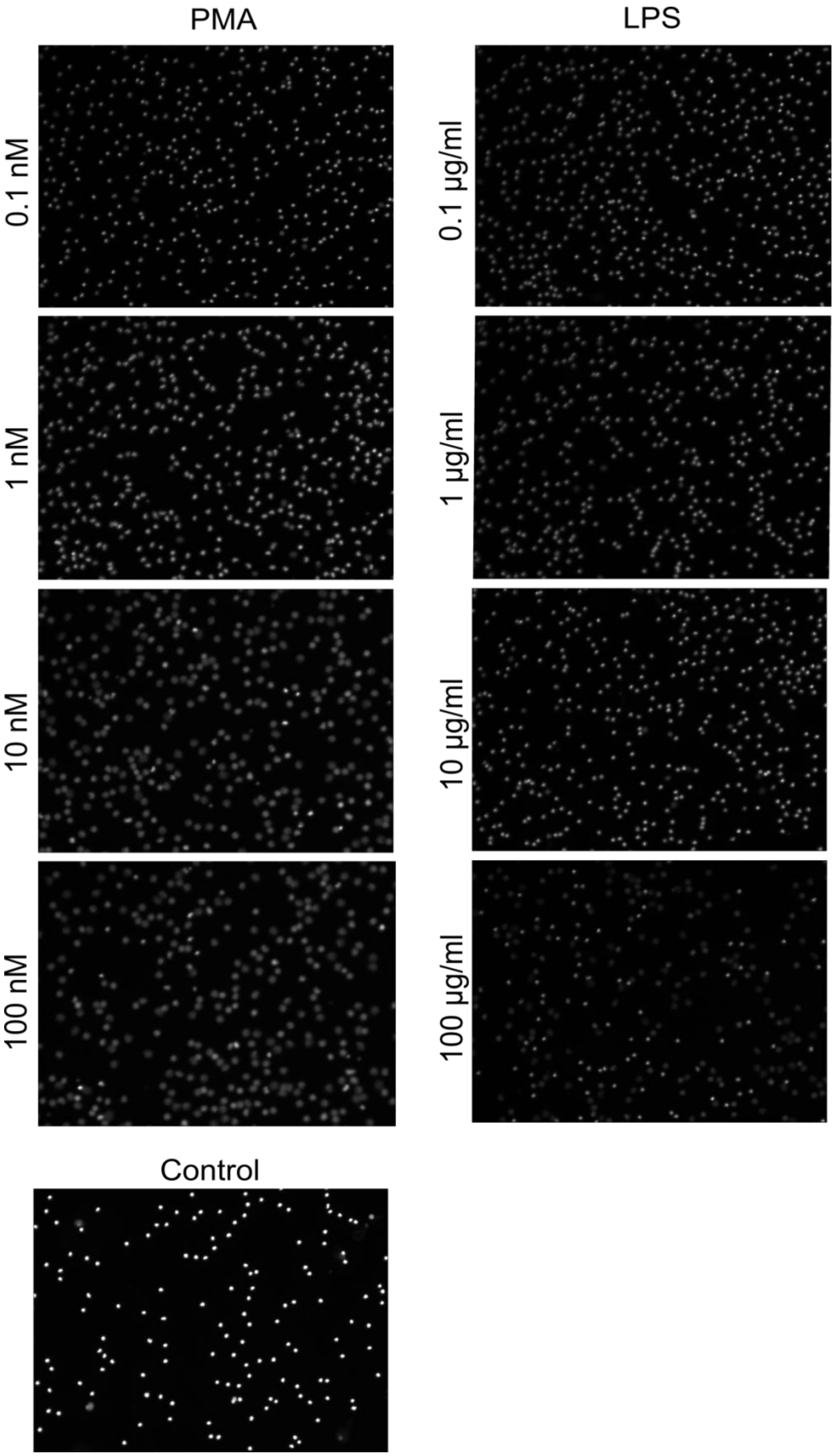

**Suppl. Fig. S4 Decondensation/NETosis behavior of activated neutrophils after 160 minutes.** Both, lower amounts of LPS and PMA (*i.e.* 0.1 – 1nM PMA and 0.1 – 1µg/ml LPS) did not show any significant decondensation compared to the control samples. In contrast, 10 – 100 nM PMA resulted in a nearly complete decondensation of all cells comparable to 100 µg/ml LPS.

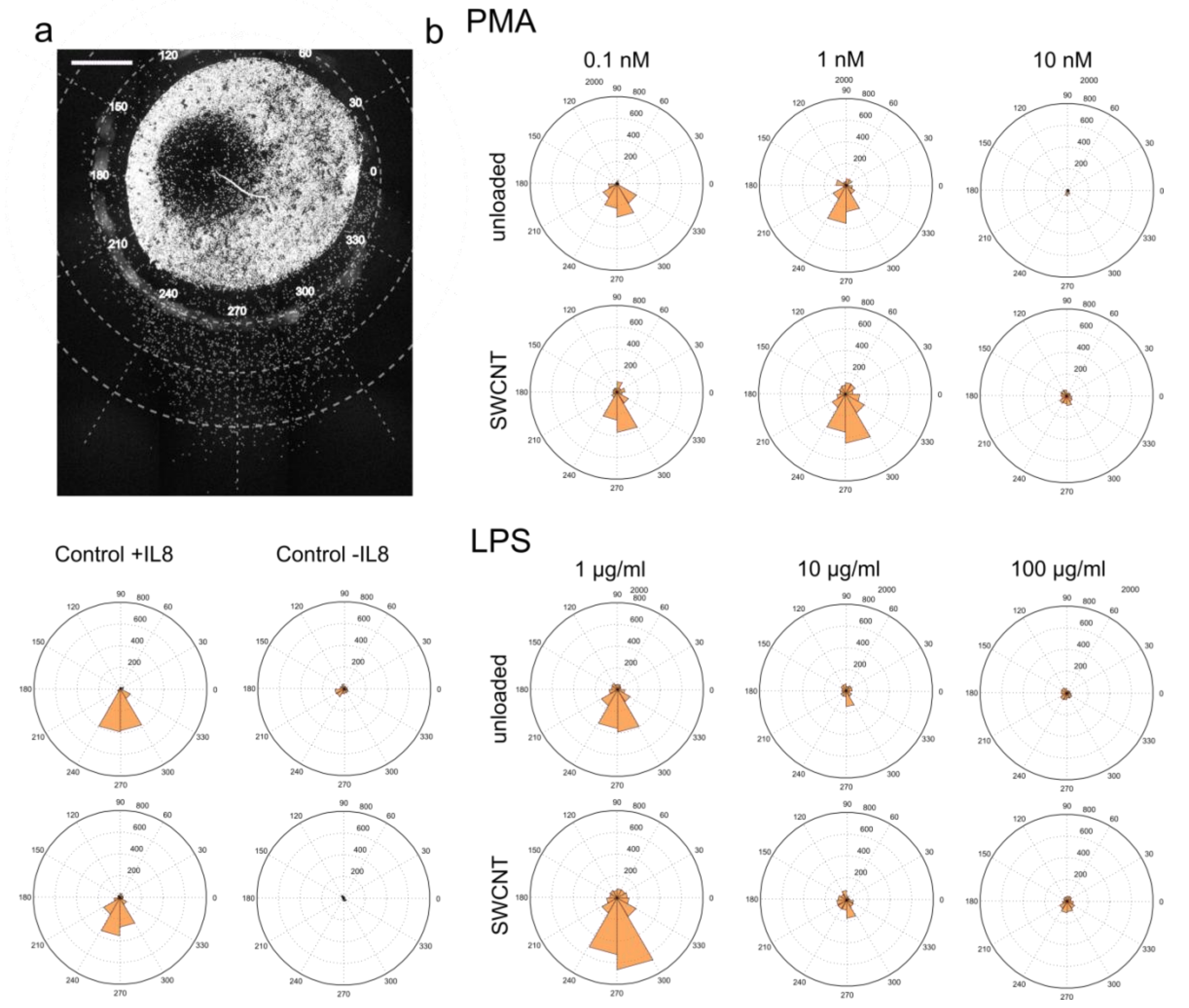

**Suppl. Fig. S5 Migration behavior of $(GT)_{15}$-loaded and unloaded neutrophils in a gradient(migration under agarose assay) NET formation inducers at 0.5% FCS concentration. a** Exemplaric image of a typical under agarose gradient experiment and the description of the coordinate system. Cells were loaded in one well (diameter around $d$ = 3 mm) and activated shortly before the chemoattractant (IL-8, 0.1 µM) was poured in a second well (distance $d$ = 2.2 mm), which generated a consistent gradient within the gel. Neutrophils were allowed to move freely for 3 hours and were imaged (Hoechst 33342 staining). Images were arranged in such a way that the 270° faces towards the IL-8 well, scale bar = 400µm. **b** Analysis of the migratory distance of $(GT)_{15}$-loaded neutrophils and untreated ones after 3 hours within an under agarose sample. Cells with SWCNTs showed enhanced movement compared to those without SWCNTs. Increased LPS or PMA concentrations reduced movement in both cases. Data was generated by measuring the maximal distance between the cell bulk and the well's edge for the respective angles. Experiments were performed three times with three independent donors (n = 3) and results were averaged to show mean values in µm (values are presented in **Suppl. Table T1**).

| **Maximal distance [µm]** | **Unloaded** | **SWCNT** |
|---|---|---|
| **PMA (0.1 nM)** | 308 ± 62 | 373 ± 104 |
| **PMA (1 nM)** | 349 ± 71 | 454 ± 109 |
| **PMA (10 nM)** | 54 ± 36 | 70 ± 15 |
| **LPS (1 µg/ml)** | 394 ± 124 | 663 ± 80 |
| **LPS (10 µg/ml)** | 151 ± 35 | 224 ± 80 |
| **LPS (100 µg/ml)** | 111 ± 20 | 145 ± 44 |
| **Control + IL8** | 395 ± 101 | 399 ± 113 |
| **Control – IL8** | 35 ± 19 | 48 ± 26 |

**Suppl. Table T1 Average distances traveled by the migrating front for 0.5% FCS conditions.** Table shows the mean distances reached by migrating front (extracted from the 255° or 285° data shown in **Suppl. Figure S5**). Distances are in µm, n = 3, Data ± SEM

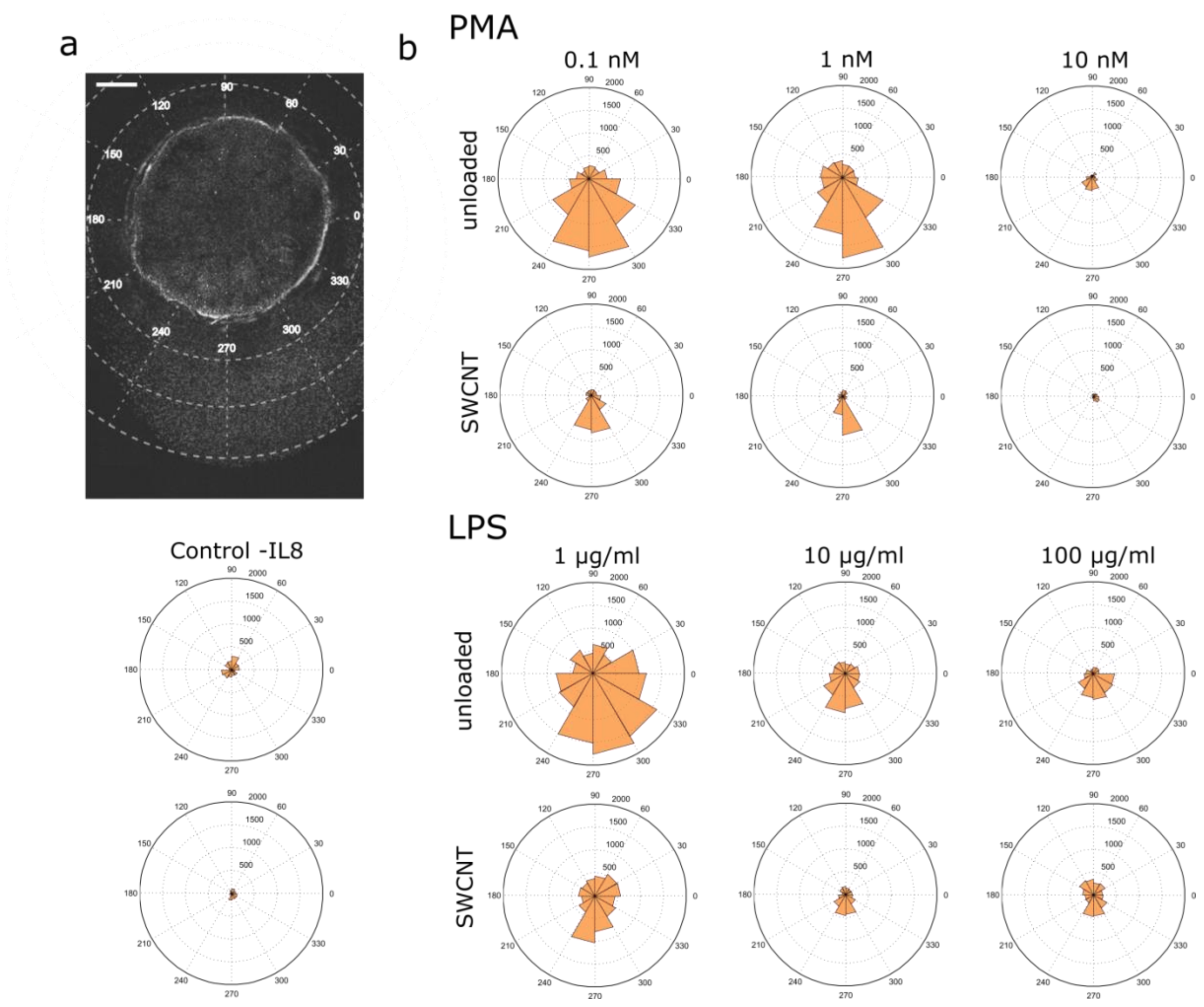

**Suppl. Fig. S6 Migration behavior of $(GT)_{15}$-loaded and unloaded neutrophils in an migration under agarose experiment using different amounts of NETosis activators and 20% FCS inherited agarose gels. a** Exemplaric image of a typical under agarose experiment plus decribtion of the used coordinate system. Cells were loaded in one well (diameter around *d* = 3 mm) and activated shortly before the chemoattractant (IL-8, 0.1 µM) was poured in a second well 2.2 mm away from the cells which generated a consistent gradient within the gel. Neutrophils were allowed to move freely for 3 hours and were imaged using Hoechst 33342 stain afterwards. Pictures were arranged so that the 270° faces towards the IL-8 well, scale bar = 500µm. **b** Analysis of the migratory distance of $(GT)_{15}$-loaded neutrophils and untreated ones after 3 hours within an under agarose sample. Cells without SWCNTs showed enhanced movement compared to those which came in contact with the sensors and even reached the other well sometimes. Increasing the concentrations of LPS or PMA resulted in reduced locomotion in both cases. Data was generated by measuring the maximal distance between the cell bulk and the well's edge for the respective angles. Furthermore, experiments were performed two times with two independent donors (n = 2) and results were averaged to show mean values in µm (exact values are presented in **Suppl. Table T2**).

| **Maximal distance [µm]** | **Unloaded** | **SWCNT** |
|---|---|---|
| **PMA (0.1 nM)** | 1724 ± 343 | 823 ± 247 |
| **PMA (1 nM)** | 1762 ± 388 | 850 ± 237 |
| **PMA (10 nM)** | 317 ± 127 | 141 ± 62 |
| **LPS (1 µg/ml)** | 1781 ± 481 | 1032 ± 235 |
| **LPS (10 µg/ml)** | 869 ± 150 | 465 ± 101 |
| **LPS (100 µg/ml)** | 578 ± 183 | 478 ± 162 |
| **Control – IL8** | 145 ± 32 | 170 ± 51 |

**Suppl. Table T2 Average distances traveled by the migrating front for 20% FCS conditions** Table shows the mean distances reached by migrating front (extracted from the 255° or 285° data shown in **Suppl. Figure S6**). Distances described in µm, n = 3, Data ± SEM

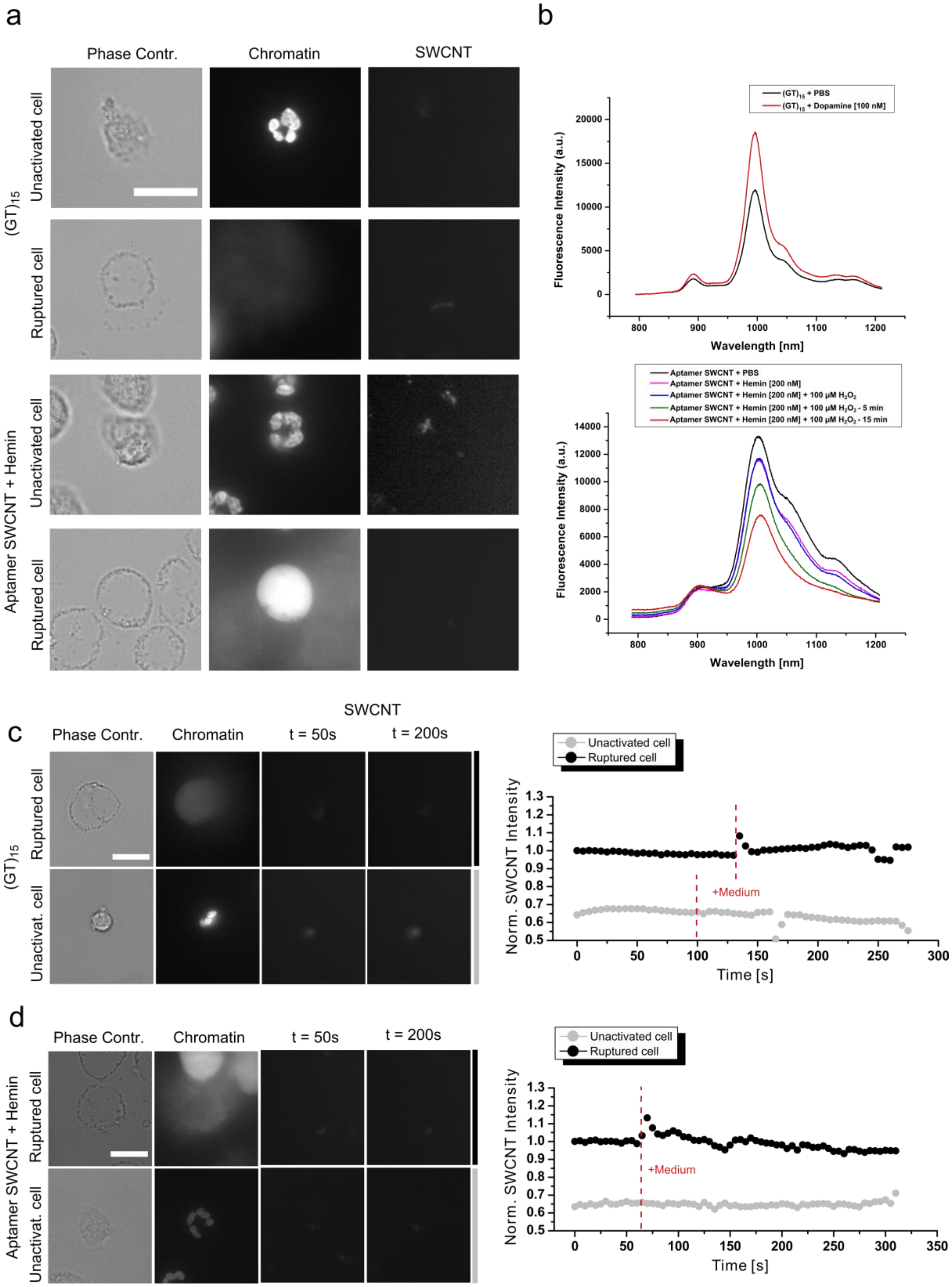

**Suppl. Fig. S7 Optical response of (GT)15- and Aptamer/Hemin SWCNTs. a** Exemplaric cell and nIR sensor images used to depict the SWCNT responses shown in **Fig. 5 c-d** (respective SWCNT movies can be seen in **Suppl. movie 6-9**). Contrast of nIR images were enhanced to show sensor geometry and position. Scale bar = 10 µm. **b** Excitation spectrum of (GT)15-SWCNT (top) and Aptamer/Hemin-SWCNT (bottom) in PBS. 100 nM dopamine increase the nIR fluorescence of (GT)15-SWCNT (dopamine sensor). 100 µM of $H_2O_2$ decreased the nIR fluorescence of Aptamer/Hemin-SWCNTs ($H_2O_2$) sensor. **c** (GT)15-SWCNTs loaded activated/non-activated cells show no significant fluorescence change when RPMI-medium is added as a control. **d** Similar control for Aptamer/Hemin SWCNT loaded activated/non-activated cells. Scale bar = 10 µm.

### Supplementary methods

#### nIR fluorescence spectroscopy

nIR fluorescence spectra were recorded with a Shamrock 193i spectrometer (Andor Technology Ltd., Belfast, Northern Ireland) connected to an IX53 microscope (Olympus, Tokyo, Japan). Excitation was performed with a 561 nm Cobolt Jive™ laser (Cobolt AB, Solna, Sweden) for $(GT)_{15}$-SWCNT samples or with an 785 nm iBeam smart laser (Toptica Photonics, Germany, Munich) in case of Aptamer-SWCNTs.

To test $(GT)_{15}$-SWCNTs responses to dopamine, 180 µl of a 0.1 nM ssDNA/SWCNT solution in PBS was placed in a 96-well plate, measured and 20 µl of 1 µM dopamine was added subsequently to yield a final concentration of 100 nM. The fluorescence response of the Aptamer/Hemin-SWCNTs was studied by adding 20 µl of 1 mM $H_2O_2$ solution in PBS to 180 µl of 2 nM Aptamer/Hemin-SWCNT with 200 nM hemin in it.
